# Multiple particle tracking detects changes in brain extracellular matrix structure and predicts neurodevelopmental age

**DOI:** 10.1101/2020.04.20.050112

**Authors:** Michael McKenna, David Shackelford, Hugo Ferreira Pontes, Brendan Ball, Tora Gao, Elizabeth Nance

## Abstract

Brain extracellular matrix (ECM) structure mediates many aspects of neuronal function. Probing changes in ECM structure could provide insights into aging and neurological disease. Herein, we demonstrate the ability to characterize changes in brain ECM structure using multiple particle tracking (MPT). MPT was carried out in organotypic rat brain slices to detect induced and naturally occurring changes in ECM structure. Induced degradation of neural ECM led to a significant increase in nanoparticle diffusive ability in the brain extracellular space. For structural changes that occur naturally during development, an inverse relationship existed between age and nanoparticle diffusion. Using the age-dependent dataset, we applied extreme gradient boosting (XGBoost) to generate models capable of classifying nanoparticle trajectories. Collectively, this work demonstrates the utility of MPT combined with machine learning for measuring changes in brain ECM structure and predicting associated complex features such as developmental age.

## Introduction

The extracellular spaces (ECS) of brain tissue are home to the brain extracellular matrix (ECM), a heterogeneous collection of proteoglycans, tenascins, and a hyaluronic acid backbone that can either be free floating, tethered to cellular surfaces, or condensed to form specific structures (Krishnaswamy et al., 2019; Zimmermann and Dours-Zimmermann, 2008). The ability to organize into specific structures allows ECM to perform unique functions that help maintain normal brain function. For example, the formation of highly condensed perineuronal nets (PNNs), which envelop the soma of certain populations of neurons in the brain, helps regulate plasticity and protects neurons from harmful processes like excitotoxicity and oxidative stress (Cabungcal et al., 2013; Okamoto et al., 1994). The basement membrane, which consists of proteoglycans, laminin, and collagen, is a three-dimensional structure that wraps around brain endothelial cells and regulates the blood-brain-barrier (BBB) and the neurovascular unit (Thomsen et al., 2017; Xu et al., 2019). Brain ECM is also highly dynamic, and the ability to assemble, disband, and reorganize is required for the development of proper neuronal circuitry and helps facilitate repair in response to injury (Barritt et al., 2006; Carstens et al., 2016; Carulli et al., 2010; Massey et al., 2006; Pizzorusso et al., 2002; Sorg et al., 2016). The structural integrity of PNNs is thought to be impacted by many neurological diseases, including epilepsy, schizophrenia, and stroke (Dzyubenko et al., 2018; Pantazopoulos and Berretta, 2016; Sorg et al., 2016; Wen et al., 2018). However, probing real-time changes in ECM microstructure, particularly changes that occur locally at the cellular level in living tissue, remains an ongoing challenge. This prevents a complete understanding of the role disease-induced changes in ECM structure play in impairing neuronal function.

To address this knowledge gap, we want to characterize changes in ECM structure that occur both spatially and temporally. This requires a technique that can probe extracellular dynamics in real-time at the microscale. Electron microscopy has been used to quantify ECS-related parameters. However, Korogod *et al*. showed that chemical fixation results in significantly smaller estimates of ECS volume fraction compared to cryo-fixation and reduces the volume of the cortex by 31% (Korogod et al., 2015). Fluorescent-based staining can be used to label specific components of brain ECM, but features commonly quantified from fluorescent images, such as fluorescence intensity and stain area, provide no direct insight into physical and geometric properties of the local environment like viscosity and ECM pore size (Lipachev et al., 2019; Rowlands et al., 2018). Atomic force microscopy (AFM) has also been used to quantify mechanical properties of brain ECM (Moeendarbary et al., 2017). However, AFM only provides a surface-level analysis, preventing analysis of the microrheological properties in the bulk of the tissue.

Multiple particle tracking (MPT) is a technique that leverages fluorescent microscopy to capture the motion of nanoparticles in real-time. MPT is unique in that the microscopic behavior of hundreds to thousands of individual particles can be tracked simultaneously, while retaining single particle resolution. The motions exhibited by particles provide information about the environment in which the particles reside, and the ability to track the movement of individual particles provides high spatial resolution. This phenomenon has already been leveraged to characterize structural features of many biological domains, including the vitreous of the eye (Xu et al., 2013), various mucosal membranes (Lai et al., 2007; Macierzanka et al., 2014; Suk et al., 2009; Wang et al., 2008), and intracellular environments (Suh et al., 2003; Suk et al., 2007; Xiao and Samulski, 2012). In the brain specifically, MPT as well as single nanoparticle tracking have been used to better estimate the width of ECS (Godin et al., 2017; Nance et al., 2012) and evaluate the diffusive ability of many nanoparticle-based drug delivery platforms (Joseph et al., 2018; Nance et al., 2014; Nance et al., 2012). An additional advantage of MPT is the sheer amount of data it generates, with experiments typically producing anywhere between 10^2^ and 10^5^ total trajectories. Because of this, machine learning methods are becoming incorporated into the MPT workflow to explore otherwise hidden trends in data and make predictions. The utility of this approach is already well documented. Wagner *et al*. demonstrated the ability to predict motion type (confined, directed, anomalous, normal) using random forest classifiers trained on trajectory feature datasets (Wagner et al., 2017), and others have employed artificial neural networks to predict agarose gel stiffness and *in vitro* cell uptake of nanoparticles (Curtis et al., 2019a).

The findings we present herein are twofold. We first demonstrate the use of MPT to characterize changes in brain ECM structure, then implement extreme gradient boosting (XGBoost) to generate classifiers capable of predicting chronological age from nanoparticle trajectory features. Moving forward, MPT can be applied to probe mechanisms that give rise to structural alterations in ECM that cause aberrant neuronal function. MPT will also provide an enhanced understanding of ECM rearrangements that occur naturally during development, aging, and pathological aging. Lastly, our results show the potential for the combined approach of MPT and machine learning to be extended to develop models capable of predicting the presence and severity of neurological disease based on nanoparticle diffusion information.

## Results

### Inducing the breakdown of PNNs in rat brain tissue *ex vivo*

We first demonstrate the ability to induce ECM structural changes in rat brain tissue *ex vivo*. This was carried out to provide a test scenario to determine whether MPT could be used to detect induced changes in ECM structure. Organotypic hemispheric brain slices taken from postnatal (P) day 35 rats were treated with either Chondroitinase ABC (ChABC, 0.4 U/mL) or hyaluronidase (HYase, 35 U/mL), two enzymes known to degrade components of brain ECM (Carstens et al., 2016; Kul’chitskii et al., 2009; Sun et al., 2018). Brain slices treated with enzyme-free slice culture media (SCM) served as the negative control (non-treated, NT). We monitored the presence of PNNs following treatment by staining with a fluorescently-labeled Wisteria Floribunda Agglutinin Lectin (WFA) at 15, 30, 45, and 120 minutes post-treatment (Figure 1A). PNN structure was completely lost in the cortex within 120 min of enzyme treatment (Figure 1B). PNN structures in non-treated brain slices were unaffected over the experimental window (Figure 1B).

**Figure 1.**
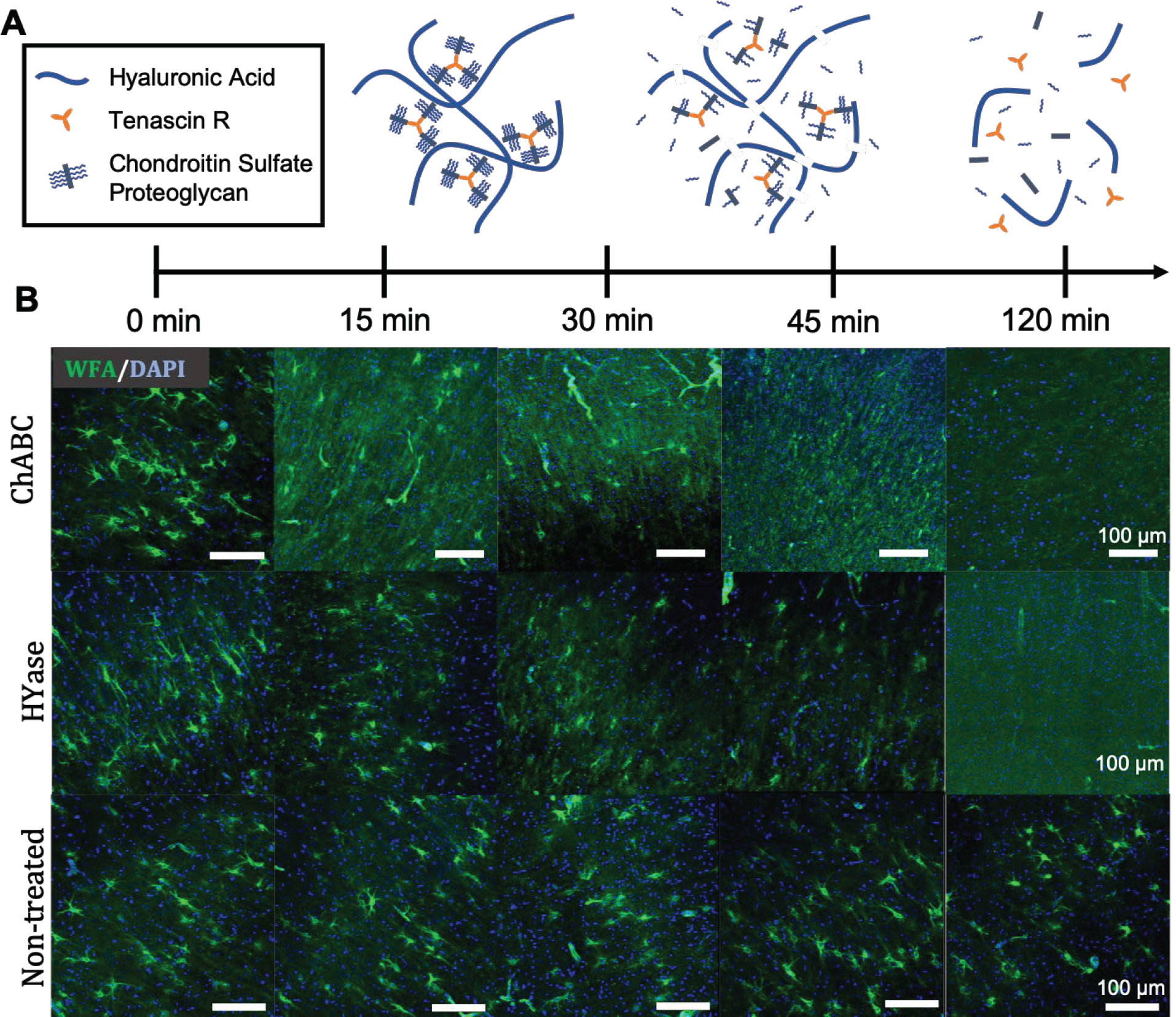
Timeline of PNN breakdown in rat brain slices *ex vivo.* (A) Schematic representation of PNN breakdown following treatment with HYase or ChABC. (B) Representative 20x magnification images taken from the cortex of P35 rat brain slices receiving one of three treatments (HYase, ChABC, or SCM). PNNs were stained with WFA (green) and cell nuclei stained with DAPI (blue). Rows represent treatment group. Columns represent treatment time. Scale bars: 100 µm.

To ensure treatment conditions did not impact brain slice viability, we monitored the release of lactate dehydrogenase (LDH) from age-matched P35 hemispheric brain slices for 23 h following treatment with identical amounts of ChABC (0.4 U/mL), HYase (35 U/mL), and SCM (non-treated, NT). LDH, an intracellular enzyme that is only released extracellularly when the cell membrane becomes compromised, provides a measure of slice viability (Su et al., 2011). Both enzyme treatment groups led to minimal cell death (<33% cytotoxicity) compared to a Triton X-100 (C_14_H_22_O(C_2_H_4_O)_10_)-treated positive control and no significant increase in cell death over the non-treated negative control (Figure S1). Collectively, these results demonstrate our ability to induce ECM structural changes in acute hemispheric brain slices while retaining slice viability.

### MPT in enzyme-treated rat brain slices *ex vivo*

Having identified the time required to cause complete loss of PNN structure in brain slices treated with either ChABC (0.4 U/mL) or HYase (35 U/mL), we next investigated whether changes in the diffusion of near neutral, poly(ethylene glycol) (PEG)-coated 40nm polystyrene nanoparticles (PS-PEG) were sensitive enough to pick up PNN structural breakdown (Table S1). We chose 40nm PS-PEG nanoparticles for two reasons. First, their 51nm hydrodynamic diameter falls below the most recently reported mean and median values of brain ECS width (Godin et al., 2017; Tonnesen et al., 2018). Second, PEG-coated PS nanoparticles have demonstrated an ability to evade adhesive interactions with various cellular and ECM-associated components (Nance et al., 2012) and remain stable in physiologically relevant conditions (Curtis et al., 2018). By evading electrostatic and hydrophobic interactions while remaining colloidally stable, the motion these particles exhibit is predominantly impacted by local fluid properties of the brain ECS and structural properties of the local ECM.

MPT in the cortex of hemispheric brain slices revealed that nanoparticle populations explore a greater area, move faster, and have increased diffusivities when diffusing in enzyme-treated brain tissue. Representative trajectory maps generated from a single video taken in a NT, ChABC-treated, and HYase-treated brain slice are provided in Figure 2A. Despite each map containing around the same number of total trajectories (1478, 1593, and 1732 for NT, ChABC-treated, and HYase-treated, respectively), nanoparticles surveyed a greater fraction of the ECS when diffusing in slices treated with ECM-degrading enzymes. In enzyme-treated slices, geometrically averaged mean-squared displacements (<MSD>) of nanoparticle trajectories were greater in magnitude for all lag times between 0 and 2 s. HYase- and ChABC-treated slices were, on average, 0.80- and 0.93-µm^2^ greater than non-treated slices, respectively (Figure 2B). The Einstein-Smoluchowski relation was used to calculate an effective diffusion coefficient, D_eff_, at a 0.33 s lag time (10 frames) for all trajectories in the dataset. The resulting range of D_eff_ values were similar for all treatment groups, but the geometric mean D_eff_ value was greater in magnitude for both enzyme-treated groups compared to the non-treated control (Figure 2C). The geometric mean D_eff_ at a 0.33 s lag time was 0.096 and 0.11 µm^2^/s for HYase- and ChABC-treated slices, respectively, compared to 0.049 µm^2^/s for non-treated slices.

**Figure 2.**
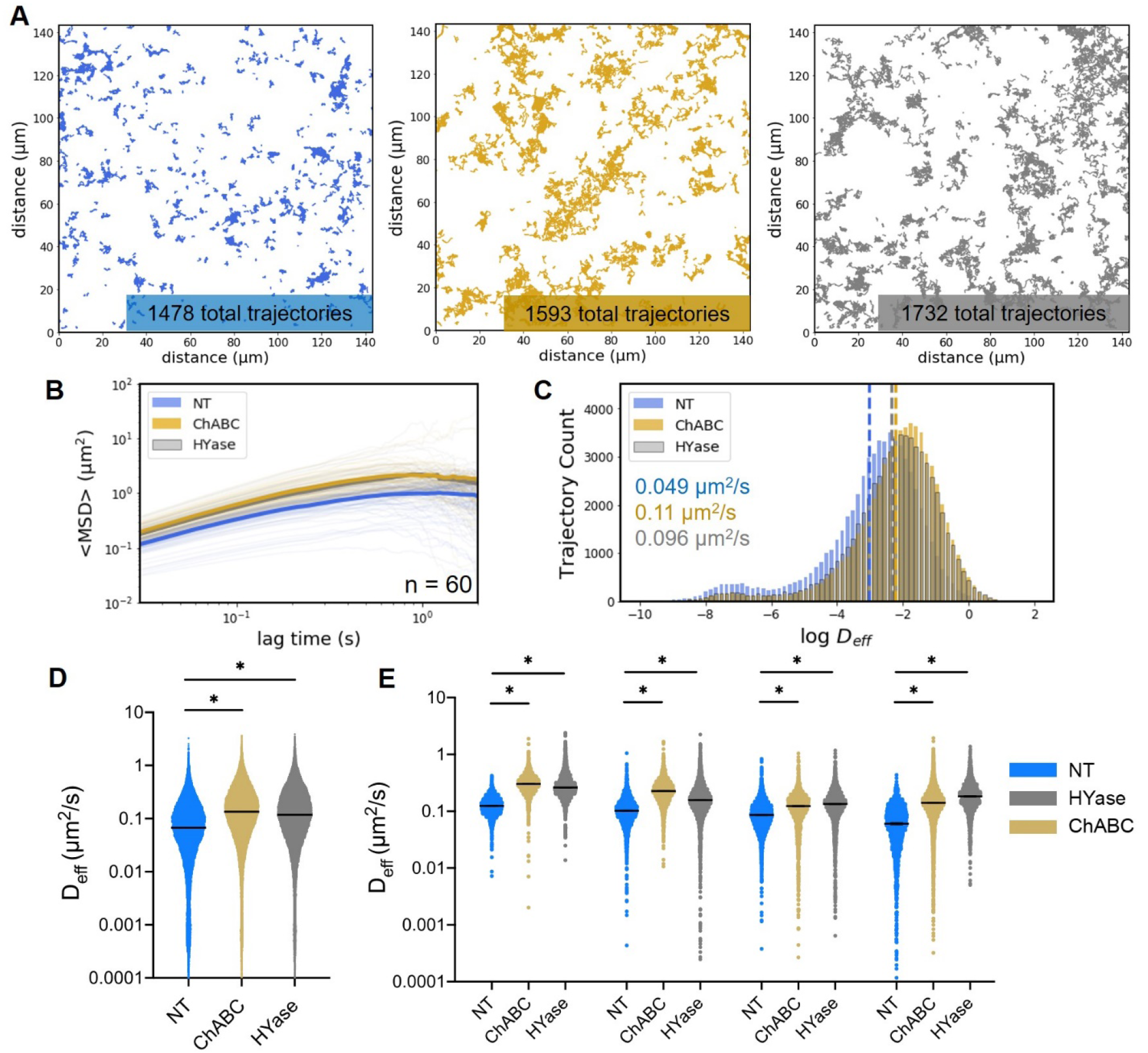
Multiple particle tracking in ChABC-, HYase-, and SCM-treated rat brain slices *ex vivo.* (A) Representative trajectory maps generated from MPT experiments carried out in non-treated (blue), ChABC-treated (gold), and HYase-treated (grey) P35 brain slices *ex vivo*. (B) Geometrically ensemble averaged <MSD> versus lag time quantified from trajectories of nanoparticles diffusing in non-treated (blue), ChABC-treated (gold), and HYase-treated (grey) P35 brain slices. Faint lines represent individual videos (60 total per group). Bold line represents the mean of 60 videos. (C) Distribution of log D_eff_ values at a 0.33 s lag time for each treatment group. The dashed lines represent the average log D_eff_ value. The geometric mean D_eff_ value is provided in writing for each treatment group. (D) Scatter plots of D_eff_ values generated from all videos (n = 5) in all slices (n = 3) in all brains (n = 4), separated by treatment group. Error bars show median value (95% CI). (E) Scatter plots of D_eff_ values from all videos (n = 5) in all slices (n = 3), separated by brain and treatment group. * denotes significant differences (Kruskal-Wallis test) between groups, after adjusting for multiple comparisons (p < 0.05).

To account for biological variability slice to slice and animal to animal, MPT experiments were performed using four separate rats, all within the same age range (P35-P38). From each animal, nine total brain slices were prepared (three for each treatment group), and five videos were collected in the cortex of each slice. Significant differences in nanoparticle diffusive ability exist independent of data grouping. If the trajectories from all videos taken in all slices and all brains are compiled into one dataset, nanoparticles diffusing in both ChABC-treated and HYase-treated slices have significantly larger D_eff_ values than those diffusing in non-treated slices (Figure 2D). If data is instead split by animal, significant differences in D_eff_ remain between treated and non-treated groups (Figure 2E). These analyses demonstrate the robustness of our findings.

To confirm particle tracking experiments did not interfere with PNN degradation, brain slices used in tracking studies were fixed at the end of the MPT window and stained with WFA and 4’,6-diamidino-2-phenylindole (DAPI) for PNNs and cell nuclei, respectively. No WFA signal was present in ChABC- and HYase-treated slices (Figure 3A). PNNs were visible in the cortex of NT slices (Figure 3A) and nanoparticles were found in close proximity to or associated with PNN structures (Figure 3B).

**Figure 3.**
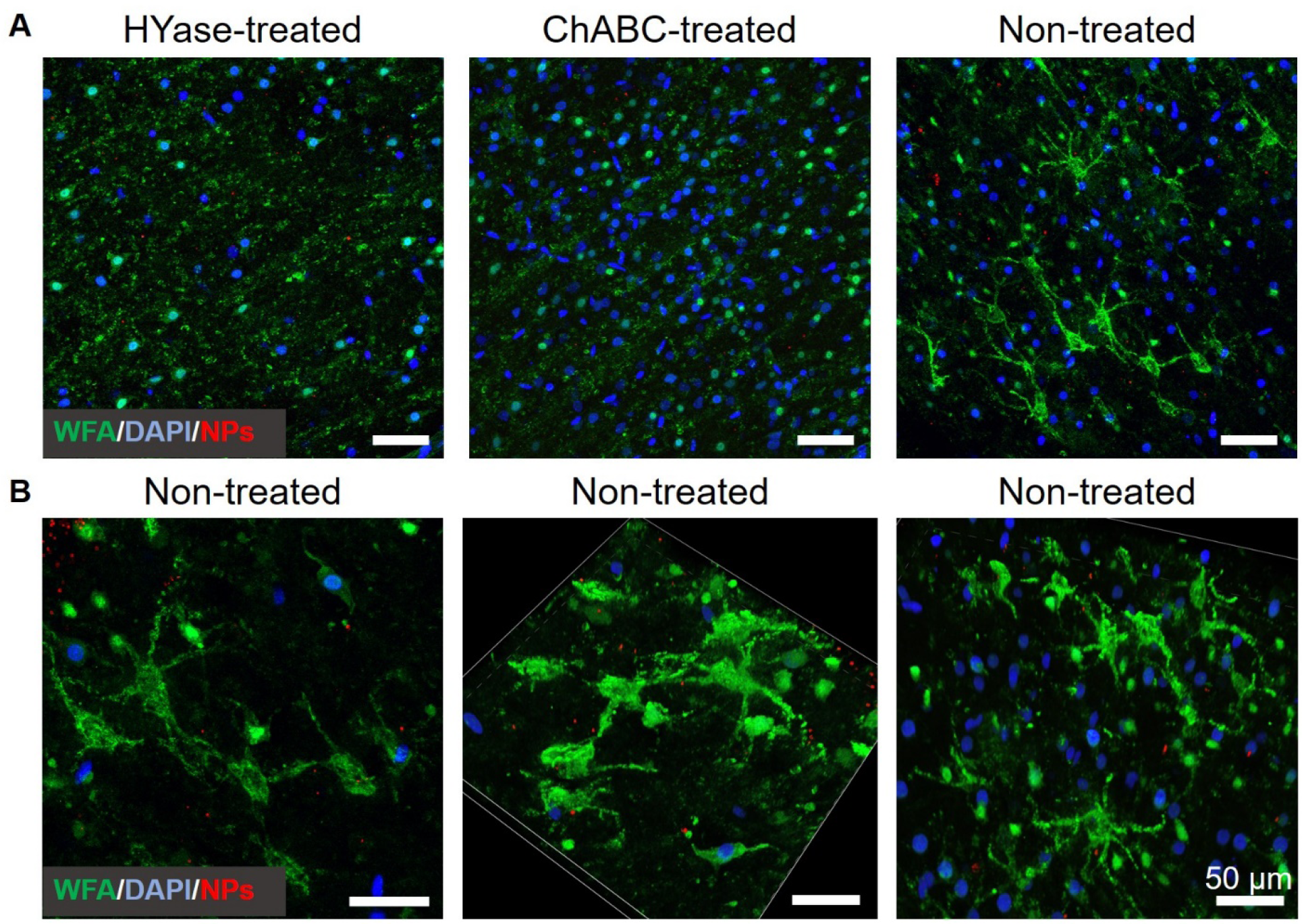
PNN imaging following MPT in treated and non-treated *ex vivo* brain slices. (A) Representative 40x magnification images taken from the cortex of P35-P38 rat brain slices post MPT. Treatment groups, represented by separate columns, were HYase, ChABC, and NT. PNNs were stained with WFA (green) and cell nuclei with DAPI (blue). Nanoparticles were red-fluorescent. (B) 60x magnification, 3D rendered z-stack images taken from the cortex of P35 rat brain slices post multiple particle tracking. All images were taken in NT slices and demonstrate the proximity of nanoparticles (red) to PNNs (green).

One potential reason for the increase in nanoparticle diffusive ability following enzymatic breakdown of ECM structures is a shift in local extracellular fluid viscosity. Both HYase and, to a lesser extent, ChABC, degrade hyaluronic acid. A shift in the distribution of hyaluronic acid molecular weights should result in a reduction of solution viscosity and subsequently reduce the amount of viscous drag experienced by the particles. To test this, solutions containing hyaluronic acid at varying molecular weights were prepared *in vitro*. The apparent viscosity (± SD) of low (33 kDa), medium (180 kDa), and high (1670 kDa) MW hyaluronic acid solutions, all at 22 mg/ml in 1xPBS, were determined to be 0.0035, 0.23, and 6.4 Pa·s, respectively (Figure 4A). MPT using 40nm PS-PEG nanoparticles showed an inverse relationship between nanoparticle diffusive ability and hyaluronic acid MW. The average D_eff_ at a lag time of 0.33 s was 0.12, 0.014, and 0.0023 µm^2^/s for particles diffusing in low, medium, and high MW hyaluronic acid solutions, respectively (Figure 4B). These *in vitro* results support our hypothesis that changes in local viscosity contribute to the observed increase in D_eff_ values following PNN degradation *ex vivo*.

**Figure 4.**
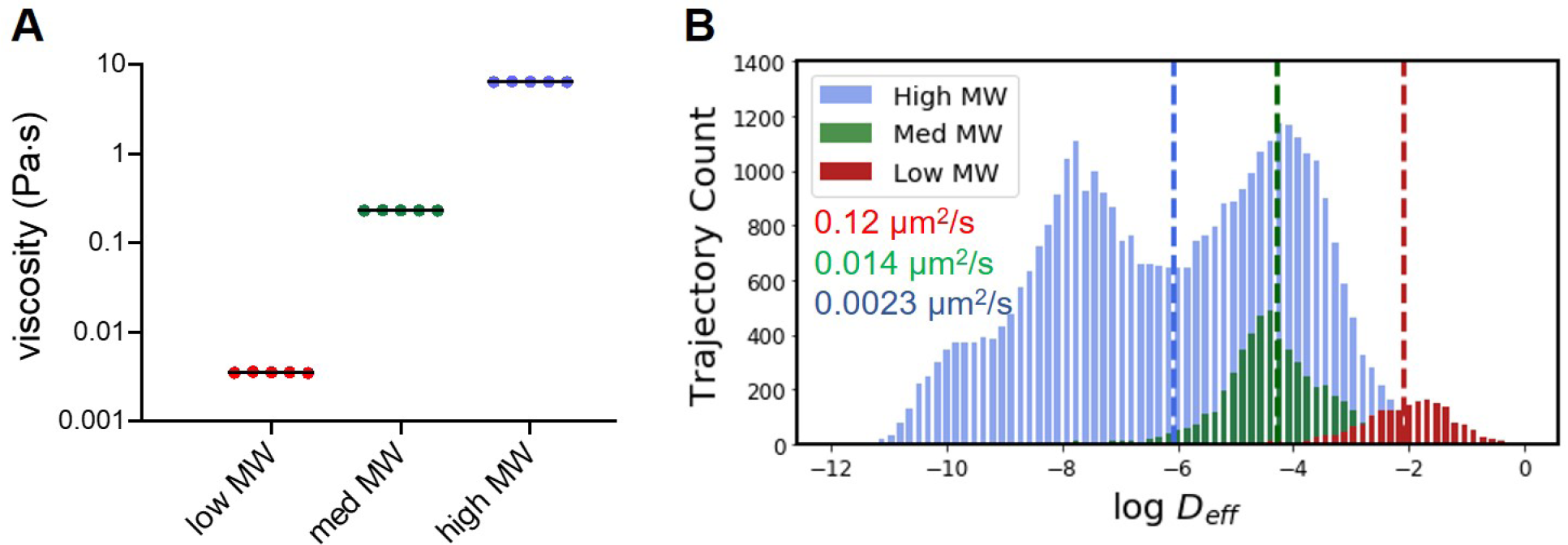
Effect of hyaluronic acid molecular weight on nanoparticle diffusive ability *in vitro*. (A) Apparent viscosity of solutions of varying hyaluronic acid molecular weight in 1x PBS (all at 22 mg/mL). Measurements (n=5) were taken at a shear rate of 100 s^-1^. (B) Distribution of log D_eff_ values at a 0.33 s lag time for each hyaluronic acid solution (data compiled from n=5 separate videos). The dashed lines represent the average log D_eff_ value. The geometric average D_eff_ value is provided in writing for each group (blue = high MW, green = medium MW, red = low MW). For all data presented, low, medium (med), and high MW correspond to 33, 180, 1670 kDa hyaluronic acid samples.

### Nanoparticle diffusive ability decreases as the density of PNNs in the cortex increases throughout development

We next show that MPT can detect naturally occurring changes in ECM structure. We leverage the brain’s tendency to form organized, function-specific ECM structures, like PNNs, during development. We first established the timeline of PNN appearance in the cortex of Sprague-Dawley (SD) rats through fluorescence-based lectin staining. PNNs stained using WFA did not appear qualitatively until 21 days after birth (P21), and preferentially form around parvalbumin-expressing (PVA^+^) interneurons (Figure 5A-D). A representative high-resolution image of a PNN taken in the cortex of a P35 rat shows the structural “mesh-like” nature of the PNN (Figure 5E). The increase in areal PNN density in the cortex between P14 and P21 was not significant, with median (95% CI) areal densities of 5.15 (3.68–6.44) and 6.32 (2.17–10.46) x10^−5^ PNNs/px^2^, respectively (Figure 5F). Significant differences did exist between P14 and P28, P14 and P35, and P21 and P35 (*p* < 0.05) groups; the median (95% CI) aerial density of PNNs in the cortex of P28 and P35 SD rats is 16.85 (13.37–22.61) and 32.85 (26.76–42.27) x10^−5^ PNNs/px^2^, respectively (Figure 5F).

**Figure 5.**
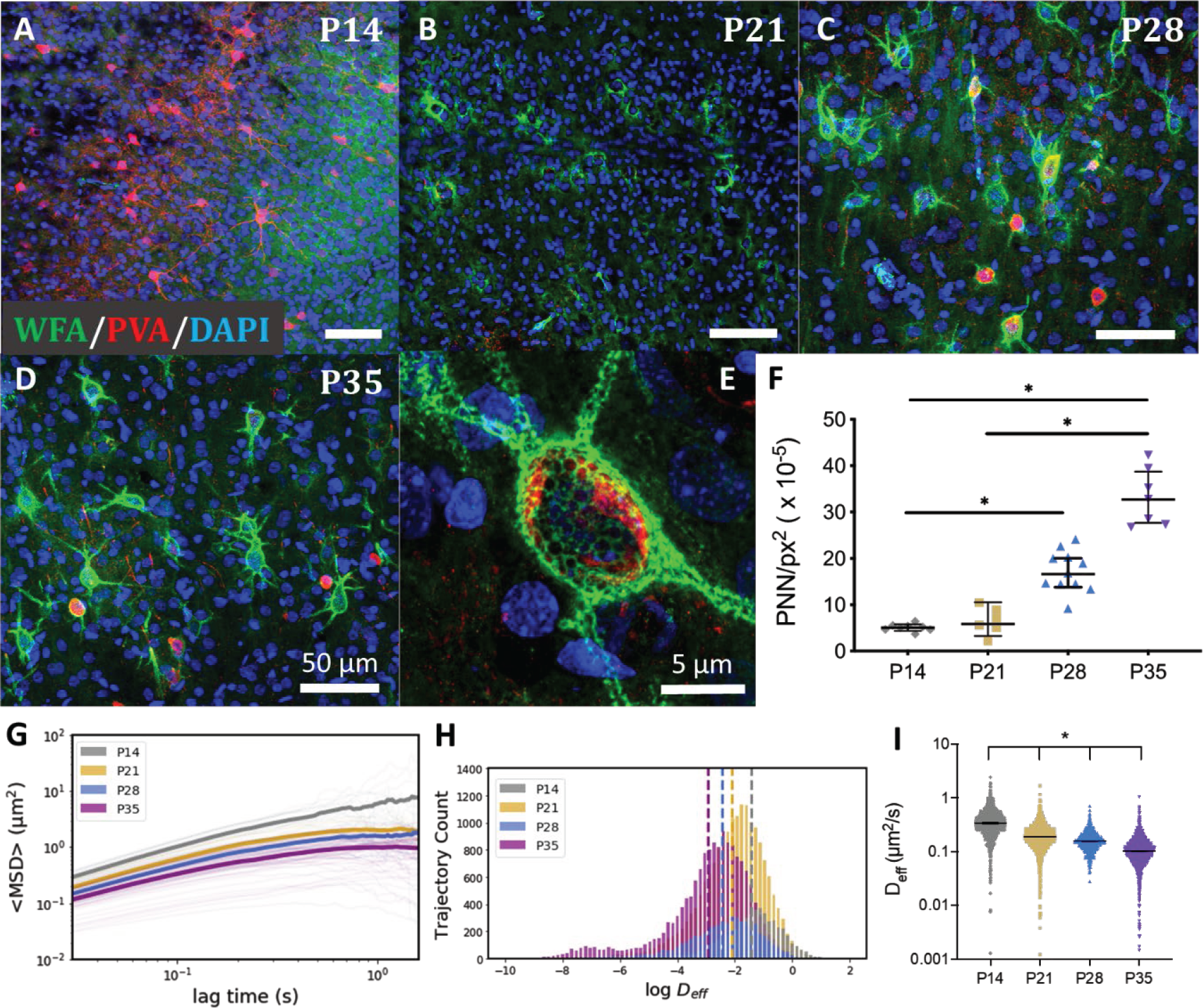
Relationship between perineuronal net formation and nanoparticle diffusion. (A-E) Representative 40x magnification images taken in the cortex of P14 (A), P21 (B), P28 (C), and P35 (D) rat brains. Brain sections were stained with WFA for PNNs (green) and anti-parvalbumin for a subpopulation of interneurons (red). Cell nuclei were stained with DAPI (blue). Scale bar = 50µm. (E) 240x magnification high resolution image of a PNN taken in the cortex of a P35 rat. Scale bar = 5µm. (F) 20x tile scans of the entire cortex were taken and the total number of PNNs was quantified. The number of PNNs per unit area increased throughout the critical period (P14-P35). Displayed are median values with a 95% confidence interval. * denotes significant differences (Kruskal-Wallis test) between groups, after adjusting for multiple comparisons. (p < 0.05). (G) Geometrically ensemble averaged <MSD> versus lag time quantified from NPs diffusing in P14 (grey), P21 (gold), P28 (blue), and P35 (purple) brain slices. Faint lines represent individual videos (15 total per group). Bold line represents the average of 15 individual videos. (H) Distribution of log D_eff_ values at a 0.33 s lag time for each pup age. The dashed lines represent the average log D_eff_ value (grey = P14, gold = P21, blue = P28, purple = P35). (I) Scatter plots of D_eff_ values generated from all videos (n = 5) in all slices (n = 3), separated by pup age. Error bars show median value (95% CI). All groups were significantly different from each other after adjusting for multiple comparisons (Kruskal-Wallis test, p < 0.05).

With the timeline of PNN formation established, MPT was performed *ex vivo* in brain slices taken from P14, P21, P28, and P35 SD rats. Tracking revealed an inverse relationship between nanoparticle diffusive ability and brain age. Geometrically averaged MSD (<MSD>) profiles decrease in magnitude as pup age increases from 14 days to 21, 28 and 35 days after birth (Figure 5G). Distributions of D_eff_ values at a 0.33 s lag time shift to lower values as the brain develops and correspondingly PNN density increases (Figure 5H). Significant differences in D_eff_ values exist between all groups (*p* = 0.05), with median (95% CI) D_eff_ of values of 0.27 (0.26– 0.28), 0.14 (0.14–0.15), 0.10 (0.10–0.11), and 0.070 (0.069–0.072) µm^2^/s for P14, P21, P28, and P35, respectively (Figure 5I).

### XGBoost classifiers provide predictive models for age-dependent MPT data

XGBoost classifiers were trained on the age-dependent MPT data to determine if the incorporation of machine learning methods could result in predictive power. Prior to model training, the amount of data the algorithm could access was enhanced by calculating additional trajectory features to complement the D_eff_ values already available. A total of 39 features were computed, some based on trajectory geometry (aspect ratio and straightness, for example) and some based on traditional diffusion theory (anomalous diffusion exponent and MSD ratio). The list of trajectory features was adopted from previous literature (Curtis et al., 2019a) but scaled up by introducing additional, local-averaged features (Table S2). An XGBoost classifier was then trained on a subset of the age-dependent feature dataset. The resulting model achieves a total predictive accuracy of 51.84% when tested on a separate subset of data, more than doubling the accuracy of a random guess (25.00%) (Figure 6). The highest predictive precision exists for the P14 and P35 groups (0.6513 and 0.5159, respectively), while the intermediate age groups (P21 and P28) are the lowest (0.4399 and 0.4110, respectively) (Figure 6A). Of the entire population of P35 trajectories included in the test dataset, only 42 were incorrectly classified as P14 (Figure 6B). Similarly, only 71 of the entire P14 population were mislabeled as P35. The majority of false predictions are contained within the middle P21 and P28 groups (Figure 6B). If we reduce the resolution of our classification by combining the P21 and P28 populations and rerun the same analysis, the model performance is elevated to an accuracy of 66.16% (Figure S2).

**Figure 6.**
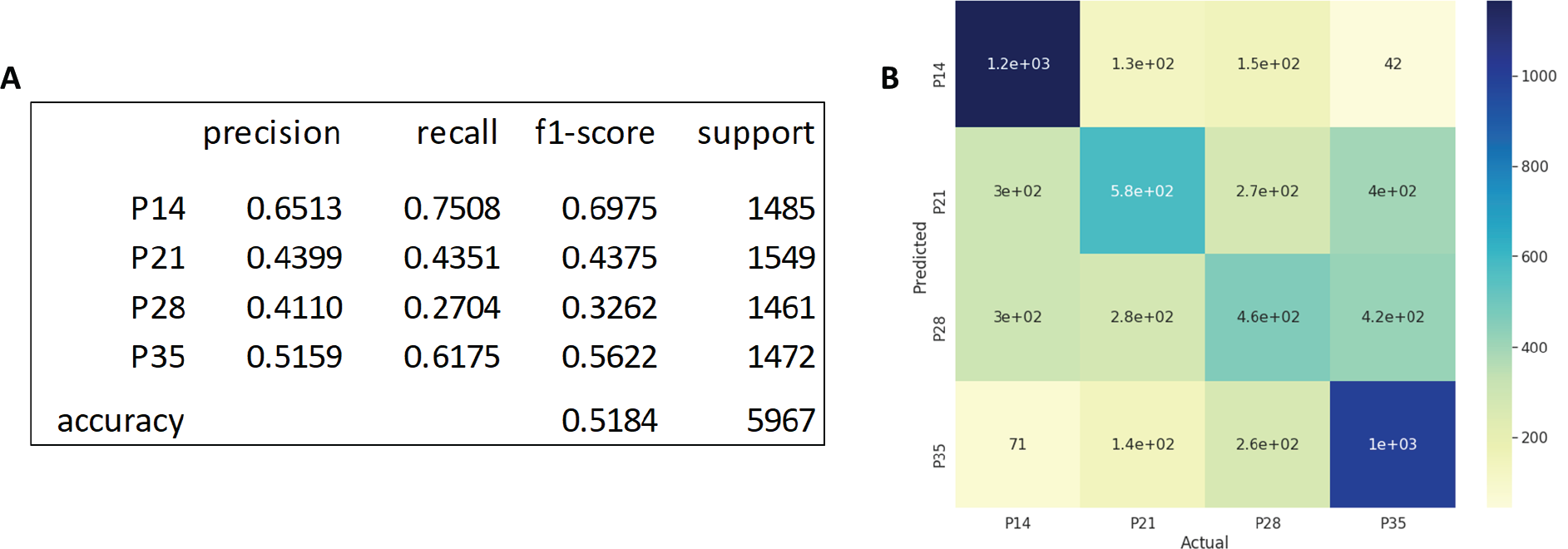
XGBoost analysis for age-dependent data. (A) Evaluation metrics for the XGBoost classifier carried out on all age groups. Included are precision, recall, f1-score, and support for each age, as well as the total accuracy and support. (B) A confusion matrix displaying how predicted outcomes compare to actual classes.

A distinct advantage of decision tree-based classifiers like XGBoost is the ability to quantify the importance of each feature to the classification being made. To identify features that influence the model most significantly, a summary plot of Shapley Additive exPlanation (SHAP) values was created for the model including all age groups (Figure 7). The top five feature dependencies are mean D_eff_ at 0.33 s (Mean Deff1), mean fractal dimension, mean fitted diffusion coefficient (D_fit), mean MSD ratio, and mean kurtosis. The least important features do not appear on the SHAP summary plot, but were determined to be trajectory elongation, trappedness, and asymmetry. None of these features contributed to model prediction. The five most important features went unchanged for the analysis performed with reduced resolution, when the P21 and P28 groups were combined (Figure S3).

**Figure 7.**
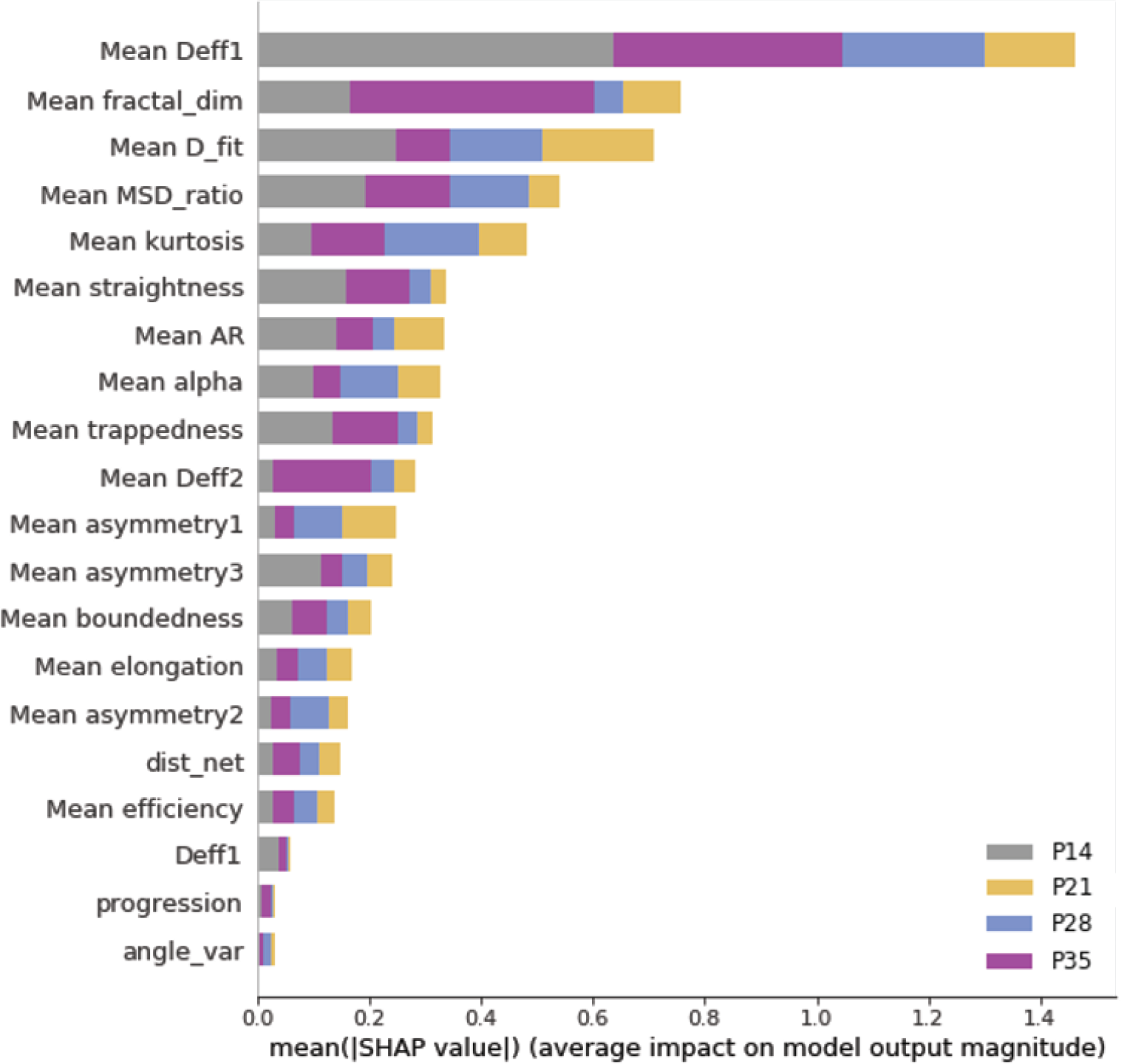
SHAP summary plot of nanoparticle trajectory features used in XGBoost classifier for age-dependent data. Provided are the 20 most influential trajectory features. Feature importance bars are color coded to provide an indication of their importance in predicting specific age groups (grey = P14, gold = P21, blue = P28, purple = P35).

Individual SHAP summary plots were created for the five most important features at each age to provide insight into how the relative importance of each feature fluctuates as both feature values and age change (Figure 8). For example, large mean Deff1 and mean D_fit values are of high importance when positively identifying the P14 group (Figure 8A). As the pups age, however, large diffusion coefficients have an inverse impact on positive prediction (Figure 8B-D). Additionally, as pups progress from 14 to 35 days old, diffusion coefficients become less important identifiers, as made evident by the reduced SHAP value and reduction in importance compared to alternative features like fractal dimension. Individual SHAP summary plots also show how the model becomes more indecisive when predicting P21 and P28 age groups. The P21 and P28 age groups contain more values that are intermediate (as indicated by purple datapoints) and closer to a SHAP value of 0.0. This provides one algorithmic-based explanation for the reduced precision of P21 and P28 groups compared to P14 and P35.

**Figure 8.**
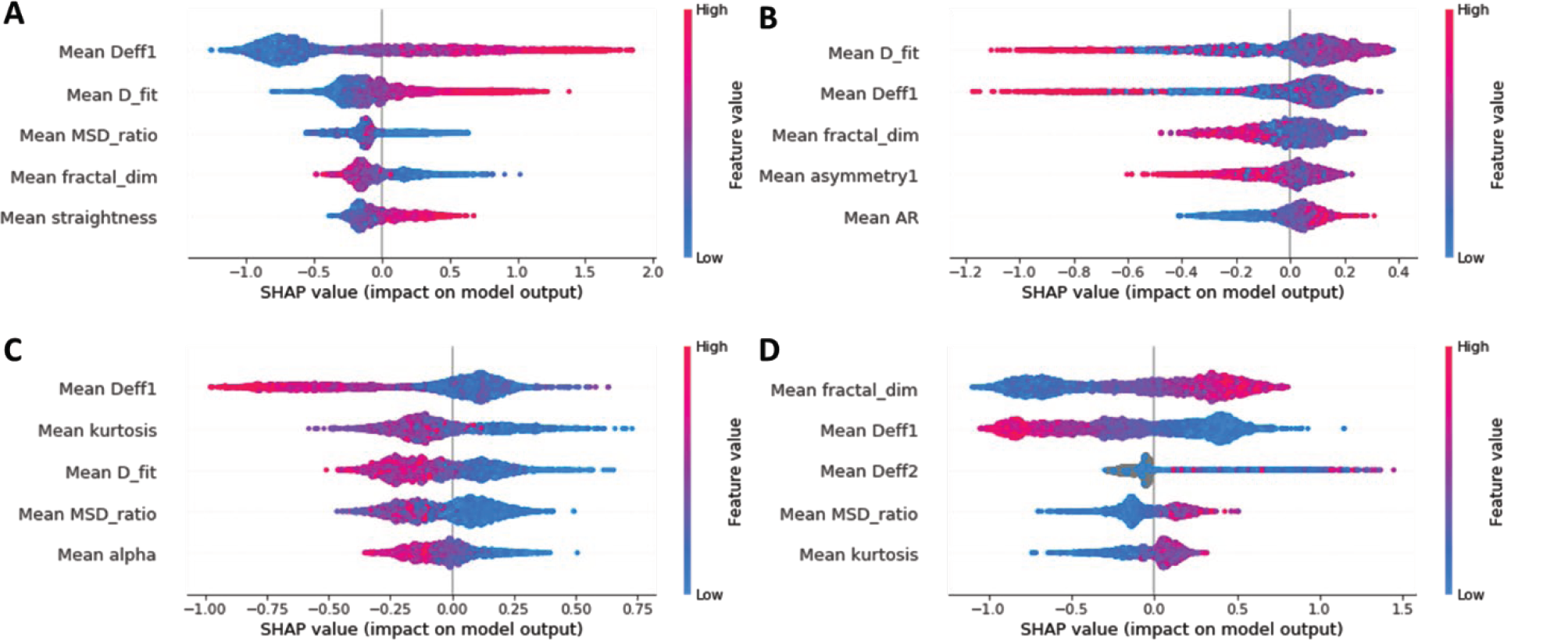
Individual SHAP summary plots for trajectory feature data. Summary plot of top five features for (A) P14, (B) P21, (C) P28, and (D) P35. Positive SHAP values further from 0.00 represent a higher impact toward positive classification in the given age category while negative SHAP values further from 0.00 represent a higher impact toward negative classification.

## Discussion

Both enzymatically induced changes in ECM structure *ex vivo* (Figure 2) and structural changes that occur naturally during development (Figure 5) brought about significant changes in the diffusive ability of 40nm PS-PEG nanoparticles. Collectively, this shows MPT can be used to capture changes in brain ECM structure. While diffusion-based techniques for characterizing the brain microenvironment already exist, researchers have been limited to either taking macroscopic approaches to quantifying diffusion related parameters (Hrabetova and Nicholson, 2007; Nicholson et al., 1979; Nicholson and Tao, 1993; Patlak and Fenstermacher, 1975; Sykova and Nicholson, 2008) or only being able to track individual particles, separately (Godin et al., 2017; Sokoll et al., 2015). Macroscopic approaches fall short in their ability to provide microscopic spatial resolution, the resolution needed to make inferences on the cellular and local extracellular level. Single particle tracking techniques provide high spatial resolution, but typically generate a single trajectory from each video collected, which in turn requires a longer time to generate datasets large enough for statistical analyses and incorporation of machine learning techniques. MPT bridges this gap. It provides a large enough sample of individual nanoparticle trajectories to make ensemble-level conclusions while also retaining high spatial resolution. This technique represents an improvement over current methods and should enhance our ability to probe structural aspects of the brain that have functional meaning, such as in the case of PNNs.

An additional advantage of MPT is that it can be readily applied to biological samples taken from other tissues and species, and even translated to *in vivo* environments, if access to proper instrumentation exists (Nance et al., 2012; Verkman, 2013). Dysregulation of ECM structure is not specific to neurological diseases. Abnormal ECM remodeling has been linked to a number of other classes of disease, including chronic pulmonary diseases (Kranenburg et al., 2006; Saetta et al., 2001), cancers (Insua-Rodriguez and Oskarsson, 2016; Sawai et al., 2008; Shields et al., 2012), and cardiovascular diseases (Fan et al., 2012; Ju and Dixon, 1996). For any and all these ailments, particularly if *ex vivo* culturing techniques exist, MPT should be considered to provide an enhanced characterization of the role ECM structural changes play in the pathological process. One relevant brain-related application would be in tissue following a traumatic brain injury (TBI). In response to TBI, glial cells near the site of injury proliferate and begin modifying the local extracellular environment to mitigate the propagation of damage and facilitate repair (Karve et al., 2016; Pekny and Pekna, 2014). Activated microglia and astrocytes release ECM degrading proteases and deposit ECM components around the injury core (George and Geller, 2018; Wang et al., 2018). The entire process of glial cell accumulation and ECM component deposition is known as glial scarring. Changes in the expression of brain ECM-specific proteoglycans in glial scars has been investigated previously, but the impact TBI has on ECM structure in both the injury core and surrounding regions remains unknown (George and Geller, 2018). MPT has the potential to better elucidate how the extracellular environment is altered in response to TBI. If targeting the injury core with a therapeutic is desired, MPT canhelp refine the design space of potential drug delivery vehicles, particularly with respect to vehicle size and surface functionalization. This represents one potential application of microstructural information generated by the MPT technique. Additional extensions of this work have potential to enhance disease diagnosis, and give insights into disease severity and disease progression.

Furthermore, we expanded the utility of biological MPT datasets by integrating a machine learning algorithm for more predictive assessments. Incorporating an XGBoost classifier into the age dependent MPT study resulted in a model capable of predicting age with 51.84% accuracy (Figure 6). Here, we limited our focus to rats grouped 7 days apart spanning a 21-day window in early postnatal development. While the model was able to predict the extreme ages, P14 and P35, with high precision, it struggled to resolve the P21 and P28 populations (Figure 6A). Recent work by Sigal et al. used a stochastic optical reconstruction microscopy technique to show that earlier in development, PNNs have greater structural heterogeneity (Sigal et al., 2019). This structural heterogeneity could necessitate larger data sets at the ages where PNNs are in the process of forming. In fact, by combining the P21 and P28 groups in one aggregate data set and retraining the model, we achieved a 66.16% accuracy (Figure S2). Despite neither classifier reaching 70% accuracy, both far exceeded the performance of a random guess, the baseline standard. Being able to accurately predict biological age to within a 1-2 day window would carry significant weight, especially if aiming to distinguish biological age from chronological age or identify the deviation from normal aging as a sign of pathological aging.

A distinct advantage of decision tree-based classifiers like XGBoost is the ability to quantify individual feature importance. Through the use of SHAP summary plots, the five most important trajectory features in accurately predicting chronological age were determined to be mean D_eff_ at 0.33 s (Mean Deff1), mean fractal dimension, mean fitted diffusion coefficient (D_fit), mean MSD ratio, and mean kurtosis (Figure 7). While the reasons why these features were most influential in predicting age falls outside the scope of this study, there is potential that certain features provide insight into specific interactions the nanoparticle is experiencing in the biological environment. For example, a shift in trajectory boundedness could represent cellular uptake, or an increase in efficiency could be indicative of particles being actively transported across a membrane (Hofling and Franosch, 2013; Wagner et al., 2017). Further work is needed to address these questions and requires the use of more controlled environments like *in vitro* cell culture. Additionally, there exist different algorithms that can be used for multiclass classification. Artificial neural networks represent a promising alternative, as they have already displayed an ability to accurately predict both nanoparticle properties like size and surface functionality, and environmental properties like gel stiffness and *in vitro* cell uptake status, when trained on trajectory feature datasets (Curtis et al., 2019a). Random forests, a form of ensemble decision trees, are another promising algorithm for classifications that exist along a continuum, having recently been applied to classifying neuroimaging data from Alzheimer’s Disease patients (Sarica et al., 2017). Therefore, the combined approach of MPT and machine learning taken herein would benefit from additional studies to determine the most optimal algorithm for this specific application. Any improvements in analytical performance and interpretability of an algorithm would accelerate progression towards predicting biological phenomena with higher granularity.

The *in vitro* work performed in solutions of varying hyaluronic acid MW elucidated one possible contributing factor to the observed changes in nanoparticle diffusive ability following a restructuring of brain ECM. *In vivo*, hyaluronic acid is produced at the cell surface and extruded through the cellular membrane into the ECS, where it acts as the main scaffolding for brain-specific ECM structures (Spicer and Nguyen, 1999; Weigel et al., 1997). Hyaluronic acid is a large, anionic, unbranched glycosaminoglycan that can reach molecular weights up to 10^7^ Da in native brain tissue (Bignami et al., 1993; Sherman et al., 2015). Given the highly anionic nature and large molecular weight of hyaluronic acid, the effect its presence can have on the movement of extracellular substances is clear. In addition to acting as a steric barrier to diffusion, diffusing substances are also subject to electrostatic interactions and local viscosity changes brought about by hyaluronic acid presence. The application of either HYase or ChABC, enzymes which degrade hyaluronic acid, to a hyaluronic acid-containing tissue sample will decrease the average MW of hyaluronic acid, reduce the local viscosity, and in turn increase the diffusive ability of extracellular substances. This phenomenon was demonstrated successfully in our *in vitro* MPT experiments, where an inverse relationship existed between hyaluronic acid MW and geometric mean D_eff_.

While a reduction in interstitial viscosity provides one possible explanation for the changes observed in *ex vivo* brain slice studies, there could be additional factors that contribute to changes in local diffusivity. Nanoparticles are subject to steric and adhesive interactions with ECM constituents and cellular surfaces, as well as hydrodynamic interactions brought about by the narrow confines of the ECS. The physical properties of the ECM are strongly dictated by the density of the biopolymers that make up the ECM, which influence the effective ECM mesh size (Engin et al., 2017). In the brain, the effect of steric constraints brought about by the ECS and ECM has been demonstrated previously. Nance et al. found D_eff_ of non-adhesive nanoparticles ranging from 40 – 200 nm decreased dramatically when nanoparticle size exceeds a certain threshold, *in vivo* in mice, and *ex vivo* in human and rat tissue (Nance et al., 2012). In reconstituted ECM systems, nanoparticles with a diameter larger than the average size of the ECM mesh are unable to penetrate, while particles smaller than the cutoff pass through (Lieleg et al., 2009). Therefore, when ECM structure condenses, as is the case for PNN formation with aging (Hensch, 2005; Testa et al., 2019), particles that were previously diffusive could experience reduced or restricted movement. This phenomenon has been demonstrated *in vitro* for biomolecular diffusion around cells embedded in a collagen gel. Kihara et al. show collagen condensation around cells results in a decrease in diffusion coefficient compared to diffusion in cell-free collagen regions, and this effect was more significant for large molecules (Kihara et al., 2013).

By reducing the size of the nanoparticle probe further than 40 nm, the size used in this study, the behavior of the nanoparticle becomes even more dependent on local ECM structure, composition, and charge distribution, and less influenced by the spatial confines of the ECS. The enhanced sensitivity to changes in ECM could give rise to more notable differences in trajectory features across different age groups, leading to more accurate predictors. However, nanoparticles with diameters smaller than the ECM mesh size can be influenced by adhesive interactions brought on by charge-charge, hydrophobic, or hydrogen bonding. The ECM network serves as a charge-selective filter, with localized charge patches (Lieleg et al., 2009). Both Nance et al. and Curtis et al. found significant differences in extracellular diffusion depending on whether PS nanoparticles had a PEG or carboxylate surface coating (Curtis et al., 2018; Nance et al., 2012). Additional studies in non-brain ECM have demonstrated the interaction between amine-modified particles with ECM protein fibrils. Researchers such as Lieleg et al. have shown that liposomal and polymer particles that are strongly charged either negatively or positively are equally unlikely to diffuse, independent of size (Lieleg et al., 2009). They further demonstrate the recovery of smaller particle diffusion when the particle charges are shielded. While this study was performed in reconstituted mouse basement membrane ECM, it’s an important insight into the impact charge-charge interactions can have on nanoparticle diffusion in the ECM. Although our study utilized densely PEG-coated particles that do not interact with the ECM (Nance, 2017), the use of charged particles to direct nanoparticle-ECM interaction could further elucidate mechanisms of ECM influence on nanoparticle diffusion in the brain ECS.

## Conclusion

The ECM plays many critical roles to maintain homeostasis in the brain. Altered ECM structure is thought to be involved in the pathophysiology of many neurological diseases, and the implications of altered PNN integrity on neuronal plasticity and activity have garnered significant attention in recent decades. In this study, we demonstrated that MPT is sensitive enough to detect changes in brain ECM structure. In addition to leveraging the brain’s natural tendency to restructure during the critical period of development, we also applied MPT to *ex vivo* hemispheric brain slices undergoing an enzymatically-induced breakdown of ECM. By incorporating XGBoost classifiers into our analysis workflow, we demonstrated the ability to use MPT data to predict chronological age. The further application of MPT in studying ECM structure could more explicitly define mechanisms involved in neurological disease progression and open new avenues of therapeutic intervention. Additionally, MPT can enhance our baseline understanding of the structure-function relationships of the brain under normal physiological conditions and has the potential to become used as one marker of neurological disease severity.

## Materials and Methods

### Organotypic hemispheric brain slice preparation

Brain slices were prepared from male SD rat pups at varying ages, depending on the specific study. This study was performed in strict accordance with the recommendations in the Guide for the Care and Use of Laboratory Animals of the National Institutes of Health (NIH). All of the animals were handled according to approved institutional animal care and use committee (IACUC) protocols (#4383-02) of the University of Washington. The University of Washington has an approved Animal Welfare Assurance (#A3464-01) on file with the NIH Office of Laboratory Animal Welfare (OLAW), is registered with the United States Department of Agriculture (USDA, certificate #91-R-0001), and is accredited by AAALAC International. Every effort was made to minimize suffering. Following euthanasia, brains were extracted, immersed in room temperature (22°C) dissection media, and cut into hemispheres with a razor blade. 300 µm-thick coronal slices were prepared from each hemisphere using a Mcllwain tissue chopper (Ted Pella, Redding, CA) (Curtis et al., 2019a). Briefly, individual slices were plated on 30 mm cell culture inserts in non-treated 6-well plates. Prior to plating, 6-well plates were filled with 1 mL SCM. Slices were incubated in sterile conditions at 37°C and 5% CO_2_. For a more detailed, step-by-step procedure of slice preparation, and buffer recipes, refer to SI Appendix, SI Experimental Procedures.

### Characterizing the timeline of enzyme-induced perineuronal net breakdown in organotypic rat brain slices *ex vivo*

All experiments were carried out within 24 h of slice preparation and used litter-matched male rats to reduce biological variability. Slices were treated with either ChABC (0.4 U/mL), HYase (35 U/mL), or SCM (NT). At the initial timepoint, 200 µL of a given treatment was applied to each brain slice and returned to the incubator. One brain slice from each treatment group was removed and fixed at 15, 30, 45, and 120 min post treatment, resulting in a total of 4 slices per treatment condition. Slices were stained with 500 µL of 1x PBS containing 10 µg/mL Fluorescein-labeled WFA Lectin for 12 h at 4°C. Cell nuclei were stained with 1 µg/mL DAPI for 30 min. All imaging was performed using a confocal microscope (Nikon Instruments, Melville, NY). Three representative images were taken at 20x magnification from the cortex of each brain slice at each time point. For specific details on ChABC, HYase, and NT working solution preparation and staining buffers, and LDH assay to measure slice viability, see SI Appendix, SI Experimental Procedures.

### Nanoparticle preparation and characterization

40nm fluorescent carboxylate (COOH)-modified polystyrene latex (PS) nanoparticles (PS-COOH) (Fisher Scientific, Hampton, NH) were covalently modified with methoxy (MeO)-poly(ethylene glycol) (PEG)-amine (NH_2_) (5kDa MW, Creative PEG Works, Winston-Salem, NC) by carboxyl amine reaction (Nance, 2017). The hydrodynamic diameter and polydispersity index (PDI) of the resulting PEG-conjugated fluorescent nanoparticles were measured via dynamic light scattering (DLS) and the ζ-potential was measured by laser Doppler anemometry. Refer to SI Appendix, SI Experimental Procedures for more detailed reaction conditions and characterization.

### Multiple particle tracking in organotypic brain slices *ex vivo*

All MPT studies were performed within 24 h of slice preparation. Slices were imaged in a temperature-controlled incubation chamber maintained at 37°C, 5% CO_2_, and 80% humidity. 30 min prior to video acquisition, injections of 40nm PS-PEG nanoparticles diluted in 1x PBS were carried out in each slice using a 10 µL glass syringe (model 701, cemented needle, 26 gauge, Hamilton Company, Reno, NV). A total of five 0.5 µL injections were made in the cortex of each slice. For the study involving the degradation of PNNs *ex vivo*, particle injections were made 90 min after treatment was applied, and videos were collected following a 30 min incubation. In total, MPT was performed 120 min after treatment with either HYase, ChABC, or SCM.

A total of five videos were collected from the cortex of each slice. Videos were collected at 33 frames-per-second and 100x magnification for 651 frames via fluorescent microscopy using a cMOS camera (Hamamatsu Photonics, Bridgewater, NJ) mounted on a confocal microscope. Nanoparticle trajectories, trajectory MSDs, and D_eff_ were calculated via diff_classifier (https://github.com/ccurtis7/diff_classifier), a Python package developed within our group (Curtis et al., 2019b).

For enzyme induced PNN breakdown experiments, three brain slices for each treatment group (ChABC, HYase, and NT) were taken from each of the four animals used. Collecting five videos from each slice resulted in a total of 60 videos and >60,000 total trajectories per treatment group. For age-dependent MPT, a total of 15 videos were taken from three slices at each age. This resulted in >4,900 total trajectories per group.

### Rheological characterization of hyaluronic acid solutions

Low (33 kDa), medium (180 kDa), and high (1670 kDa) MW hyaluronic acid samples (R&D Systems, Boston, MA) were added to separate solutions of 1x PBS to achieve a final concentration of 22 mg/mL (2.2 wt%). A rheometer (Physica MCR 301, Anton Paar, Graz, Austria) operating in rotational mode was used to measure the apparent viscosity of each solution at a shear rate of 100 s^-1^. A 25 mm parallel plate attachment (Anton Paar) was operated at a 0.5 mm gap for all experiments. The base plate was set to 22°C 30 min prior to the experiment and held constant throughout the duration of the experimental window.

### Multiple particle tracking in hyaluronic acid solutions

MPT experiments carried out in hyaluronic acid solutions *in vitro* were performed similarly to MPT experiments in brain tissue *ex vivo*. Briefly, 40nm PS-PEG nanoparticles were added to hyaluronic acid solutions and a total of five videos were collected from each solution. Videos were collected at 33 frames-per-second and 100x magnification for 651 frames via fluorescent microscopy using a cMOS camera mounted on a confocal microscope. Nanoparticle trajectories, trajectory MSDs, and D_eff_ were calculated via diff_classifier (Curtis et al., 2019b).

### Immunohistochemistry and lectin staining on fixed rat brain slices

Following euthanasia, SD rats were perfused with sterile 1x PBS. Brains were immediately extracted and placed in 10% formalin phosphate buffer for 24 h at 4°C. Brains went through a 30% sucrose gradient to be frozen and sectioned into 30 µm-thick coronal sections using a Leica CM1950 cryostat (Leica Biosystems, Buffalo Grove, IL). Sections were first incubated with rabbit anti-Parvalbumin (anti-PVA, Abcam ab11427, Cambridge, UK) at a 1:100 dilution in 1x PBS containing 1% Triton X-100 (MilliporeSigma), 3% donkey serum (MilliporeSigma), and 10 µg/mL WFA for 6 h at room temperature (22°C). Following a wash step, a 1:500 dilution of Alexa Fluor 568-labeled donkey anti-rabbit IgG (ThermoFisher) in 1x PBS containing 1% Triton X-100 and 10 µg/mL WFA was applied to sections for 2 h. Cellular nuclei were stained with a 1 µg/mL solution of DAPI in 1x PBS for 15 min. Following a final wash, microscope slides were mounted with a glass coverslip using Wako antifade media (Vector Laboratories) and stored at −20°C until imaged. Sections were imaged using a confocal microscope. Z-stack scans of the entire coronal section were taken at 20x magnification. For a more detailed methodology, image processing and PNN quantification, see SI Appendix, SI Experimental Procedures.

### XGBoost predictive model for age related data classification

XGBoost is a type of boosted decision tree in which the algorithm builds itself sequentially using multiple weak learners until a strong learner can be produced. Every tree produced in the series is fit to a modified weighted version of the original dataset. This sequential method continues until a set number of learners has been created or until the model converges within the exponential loss function. Prediction is then made by calculating the weighted average of all produced learners (Trevor Hastie, 2009). XGBoost specifically incorporates regularization into its algorithm to control overfitting the data during training. It incorporates a unique objective function that encourages simple models and decreases variance (Patryk Orzechowski, 2018).

50,444 samples were rebalanced using under-sampling into four even sets of 6000 data points for each age classification. Training and testing datasets were randomly sampled from the age data with a training/testing split percentage of 80% to 20%. Features were chosen and calculated based on the geometry of the trajectory using feature calculation algorithms on diff_classifier (https://github.com/ccurtis7/diff_classifier). This includes asymmetry, anomalous exponent, aspect ratio, elongation, boundedness, fractal dimension, efficiency, straightness, kurtosis, and MSD ratio. Extra features were created based on the immediate surrounding data.

Mean values of each calculated feature were calculated and used in prediction. In total, 39 different features were used. A comprehensive list and description of every feature used can be found in the supplemental text (Table S2). The XGBoost model was trained using a max depth of 7, an eta of 0.005, a gamma of 5, a subsample of 0.15, and a colsample_bytree of 0.8. Following initial training, feature selection was implemented to remove features that were unimportant to prediction and to improve model performance.

To better understand the individual contribution to overall prediction, shapely additive explanations (SHAP) were calculated for every feature. SHAP is based on the theoretically optimal use of Shapley Values (Lundberg, 2017), which are a feature’s contribution to the prediction, 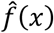:

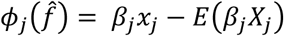

in which *Eβ*_*j*_ *X*_*j*_ is the mean effect estimate for feature j. The contribution is the difference in the feature effect and the average effect (Molnar, 2019). SHAP were used to create summary and dependency plots of the top features in prediction of each age category. The summary plot shows the average impact of each feature on prediction output calculated by the mean absolute SHAP values:

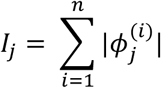

This importance value differs from other importance calculations due to its basis on magnitude of feature attributions (Molnar, 2019). Analysis for age-related data can be found on the ECM-MPT-Predictive_Age_Data repository (https://github.com/dash2927/ECM-MPT-Predictive_Age_Data).

### Statistical Analysis

All statistical analyses were carried out in GraphPad Prism (GraphPad Software Inc, Version 8.2.0). For all tests run, differences were defined as statistically significant at p < 0.05. The D’Agostino-Pearson omnibus K2 test was used to test for normality. If we were unable to reject the null hypothesis that data were sampled from a population that follows a Gaussian distribution, we ran Brown-Forsythe and Welch ANOVA tests. We used Dunnett T3 to correct for multiple comparisons. If we were able to reject the null hypothesis that the data were taken from a normally distributed population, we used the Kruskal-Wallis test for significance. In these instances, we applied Dunn’s method to correct for multiple comparisons.

## Data and Code Availability

All data presented herein can be provided upon request. All code is available on github, with links included in the methods.

## Author Contributions

M.M. and E.N. designed the experiments. M.M. conducted the experiments. D.S. developed the machine learning algorithm, trained the algorithm and performed all model validation and testing for XGBoost. M.M. and H.F.P. optimized staining protocols, performed staining and imaging of PNNs, generated slices for all ex vivo studies, and performed MPT. M.M. performed all MPT analysis. B.B. prepared hyaluronic acid hydrogels, and assisted M.M. in performing rheology and MPT experiments in gels. T.G. processed, cryosectioned, and stained brain slices different aged animals for PNN imaging. M.M. and E.N. wrote and revised the manuscript, and prepared the manuscript for submission.

## Acknowledgements

The authors would like to thank Dr. Chad Curtis, who established the Python-based analysis pipeline that was used to track and quantify nanoparticle trajectories. This work was supported by the National Institute of General Medical Sciences (Grant #1 R35 GM124677-01) and the Data Sciences Initiative in the Department of Chemical Engineering at the University of Washington.

## Supplemental Information

### SI Experimental Procedures

#### Organotypic hemispheric brain slice preparation

Animals were administered an intraperitoneal injection of pentobarbital (150 mg/kg). After euthanasia, brains were rapidly removed and immersed in room temperature (22°C) dissection media consisting of 500 mL HBSS (no Mg^2+^, no Ca^2+^, ThermoFisher, Waltham, MA), 1% Penicillin-Streptomycin (MilliporeSigma), and 3.2 g glucose (MilliporeSigma). Whole brains were cut into hemispheres with a razor blade, and 300 µm-thick coronal slices were prepared from each hemisphere using a Mcllwain tissue chopper. Slices were transferred to a Petri dish filled with room temperature dissection media and separated under a surgical dissection microscope using fine tip paint brushes. Individual slices containing corpus callosum were taken from the prefrontal cortex and placed on 30 mm cell culture inserts (Fisher Scientific) in non-treated 6-well plates (USA Scientific). Prior to plating, 6-well plates were filled with 1 mL slice culture media (SCM) containing 250 mL MEM (ThermoFisher, no glutamine, no phenol red), 125 mL HBSS (with Mg^2+^, with Ca^2+^, ThermoFisher), 125 mL horse serum (ThermoFisher), 5 mL GlutaMAX Supplement (Fisher Scientific), and 1% Penicillin-Streptomycin. Slices were incubated in sterile conditions at 37°C and 5% CO_2_.

#### Characterizing the timeline of enzyme-induced perineuronal net breakdown in organotypic rat brain slices *ex vivo*

Slices were treated with either ChABC (MilliporeSigma), HYase (Fisher Scientific), or SCM (non-treated, NT) working solution. ChABC working solution was prepared by reconstituting Chondroitinase ABC to 0.4 U/mL in an aqueous buffer containing 50mM Tris HCl, pH 8.0 (ThermoFisher Scientific), and 50mM sodium acetate (MilliporeSigma). For the HYase working solution, HYase from *Streptomyces hyalurolyticus* (Fisher Scientific) was reconstituted to 35 U/mL in 4°C 1x Dulbecco’s Phosphate-Buffered Saline (1x PBS, no Mg^2+^, no Ca^2+^, Corning). Following reconstitution, both ChABC and HYase working solutions were aliquoted and stored at −20°C until use. The NT working solution consisted of SCM.

All working solutions were brought to room temperature (22°C) prior to use. At the initial timepoint, 200 µL of a given treatment solution was applied to the top of each brain slice in a dropwise fashion and returned to the incubator, where slices were maintained at 37°C and 5% CO_2_. At subsequent timepoints, a brain slice was removed and placed in 10% formalin phosphate buffer (Fisher Scientific) for fixation. One brain slice from each treatment group was removed at 15, 30, 60, and 120 min post treatment, resulting in a total of 4 slices per treatment condition. Slices were incubated in formalin for 1 h at room temperature (22°C), washed 2 times with 500 µL 1x PBS for 5 minutes each, and stored at 4°C until staining commenced. Slices were stained within 1 week of fixation with 500 µL of 1x PBS containing 10 µg/mL WFA Lectin (Vector Laboratories Inc, Burlingame, CA) for 12 h at 4°C. Following WFA incubation, slices were washed 2 times with 500 µL 1x PBS for 5 minutes, and cell nuclei were stained with 500 µL of 1x PBS containing 1 µg/mL DAPI (ThermoFisher) for 30 min. Slices were subject to a final washing step and stored in 1x PBS at 4°C until imaged.

All imaging was performed within two weeks of staining using a confocal microscope (Nikon Instruments, Melville, NY). At a 20x magnification, 3 representative images were taken from the cortex of each brain slice.

#### Lactate dehydrogenase assay for assessment of brain slice viability

Whole hemisphere brain slice viability was evaluated using LDH assay (Cayman Chemical, Ann Arbor, MI), which measures the LDH released into the culture medium from degenerating cells in brain slices (Su et al., 2011). All experiments were carried out within 24 h of slice preparation, and all working solutions were brought to room temperature (22°C) prior to use. Two hours prior to the 0 h timepoint (−2 h), the SCM present below the membrane insert was exchanged for 1 mL of serum-free SCM consisting of 250 mL MEM (no glutamine, no phenol red), 250 mL HBSS (with Mg^2+^, with Ca^2+^), 5 mL GlutaMAX Supplement, and 1% Penicillin-Streptomycin. Immediately following media exchange (still at the −2 h timepoint), 200 µL of a given treatment solution was applied in dropwise fashion to the top of each slice. Brain slices were returned to the incubator for a 2 h treatment period, where they were maintained at 37°C and 5% CO_2_. At 0, 1, 2, 4, and 23 h after the initial 2 h treatment period, the serum-free SCM supernatant resting below the membrane insert was collected, frozen at −80°C, and replaced with 1 mL of fresh serum-free SCM that had been preheated to 37°C. For the Triton-X 100-treated positive control, no treatments were applied to the top of the slice (at the −2 h timepoint). Instead, the serum-free SCM was doped with 1% Triton-X 100 at every media exchange. The percentage of LDH released in each whole hemisphere brain slice was quantified according to manufacturer’s recommendations (Cayman Chemical). Briefly, the fluorescence intensity of the supernatant samples was measured, the background of a negative control (serum-free SCM) was subtracted, and all values were normalized to the intensity of the Triton-X 100-treated positive control (the supernatant collected at the final time point), which represented max cell death (100% cytotoxicity).

#### Nanoparticle preparation and characterization

The covalent attachment of MeO-PEG- NH_2_ (5kDa MW, Creative PEG Works) to the surface of 40nm fluorescent PS-COOH nanoparticle (Fisher Scientific) was carried out using a carboxyl amine reaction (Nance, 2017). Briefly, 50 µL of stock PS-COOH particle suspension was washed and resuspended to six-fold dilution in ultrapure water. A four-fold molar excess of MeO-PEG-NH_2_ was added to the particle suspension and mixed to dissolve the PEG. *N*-Hydroxysulfosuccinimide (NHS, MilliporeSigma, Burlington, MA) was added to a final concentration of 60 mM and 200 mM borate buffer (pH 8.2) was added to dilute the 300 µL sample volume five-fold. 1-Ethyl-3-(3-dimethylaminopropyl) carbodiimide (EDC, Invitrogen, Carlsbad, CA) was added to stoichiometrically complement the MeO-PEG-NH_2_. Tubes containing particle suspensions were wrapped in aluminum foil and placed on a rotary incubator for 6 h at 22°C and then washed via centrifugation (Amicon Ultra 0.5 mL 100k MWCO; MilliporeSigma) at conditions specified previously (Nance, 2017). Particles were resuspended in ultrapure water to the initial stock particle volume and stored at 4°C until use.

For nanoparticle characterization, both DLS and laser Doppler anemometry were performed using the Zetasizer Nano ZS (Malvern Instruments, Malvern, UK). Particles were diluted to ∼0.002% solids in filtered (0.45 um, Whatman, Maidstone, UK) 10 mM NaCl, pH 7.0, prior to measurement.

#### Immunohistochemistry and lectin staining on fixed rat brain slices

Male SD rats were first administered an intraperitoneal injection of pentobarbital (150 mg/kg). After euthanasia, rats were perfused with sterile 1x PBS. Brains were removed and immediately placed in 10% formalin phosphate buffer for 24 h at 4°C. Brains were then subjected to a sucrose gradient: 10% formalin phosphate was first exchanged for 15 weight % sucrose (MilliporeSigma) in 1x PBS and allowed to incubate for 24 h; the 15% sucrose was then exchanged for 30% sucrose in 1x PBS. Following a 24 h incubation in 30% sucrose, brains were removed from solution and frozen at −80°C until ready for use. Frozen brains were sectioned into 30 µm-thick coronal sections using a Leica CM1950 cryostat (Leica Biosystems, Buffalo Grove, IL).

Sections were first incubated in the dark with rabbit anti-PVA (Abcam ab11427, Cambridge, UK) at a 1:100 dilution in 1x PBS containing 1% Triton X-100 (MilliporeSigma), 3% donkey serum (MilliporeSigma), and 10 µg/mL WFA for 6 h at room temperature (22°C). Following primary antibody incubation, slices were washed two times for 2 min each with 1x PBS. A 1:500 dilution of Alexa Fluor 568-labeled donkey anti-rabbit IgG (ThermoFisher) in 1x PBS containing 1% Triton X-100 and 10 µg/mL WFA was then applied to the sections for 4 h at room temperature in the dark. Again, sections were washed two times with 1x PBS for 2 min. Cellular nuclei were stained with a 1 µg/mL solution of DAPI in 1x PBS for 15 min. Following a final wash step, microscope slides were mounted with a glass coverslip using Wako antifade media (Vector Laboratories) and stored at −20°C until they were imaged. Sections were imaged using a confocal microscope. Z-stack scans of the entire coronal section were taken at 20x magnification.

#### Image processing for quantifying the density of perineuronal nets in the cortex

Image processing was performed in ImageJ (Schindelin et al., 2012). First, the maximum intensity projection of the full section z-stack scan was generated and a region of interest (ROI) drawn around the entire cortex. All signal that fell outside the ROI was eliminated. The background was subtracted from the resulting image using a rolling ball radius of 5 pixels. A threshold was then applied, with the lower threshold being set to 0 and the upper threshold set by the user. The image was then subjected to a dilation, holes were filled, and a watershed applied. The total number of PNNs was quantified using the *Analyze Particles* plugin, with the minimum particle size set to 50 px^2^. The total number of PNNs was normalized to the area of the ROI. Experimental groups were separated based on brain age.

## Supplemental Figures and Tables

**Figure S1.**
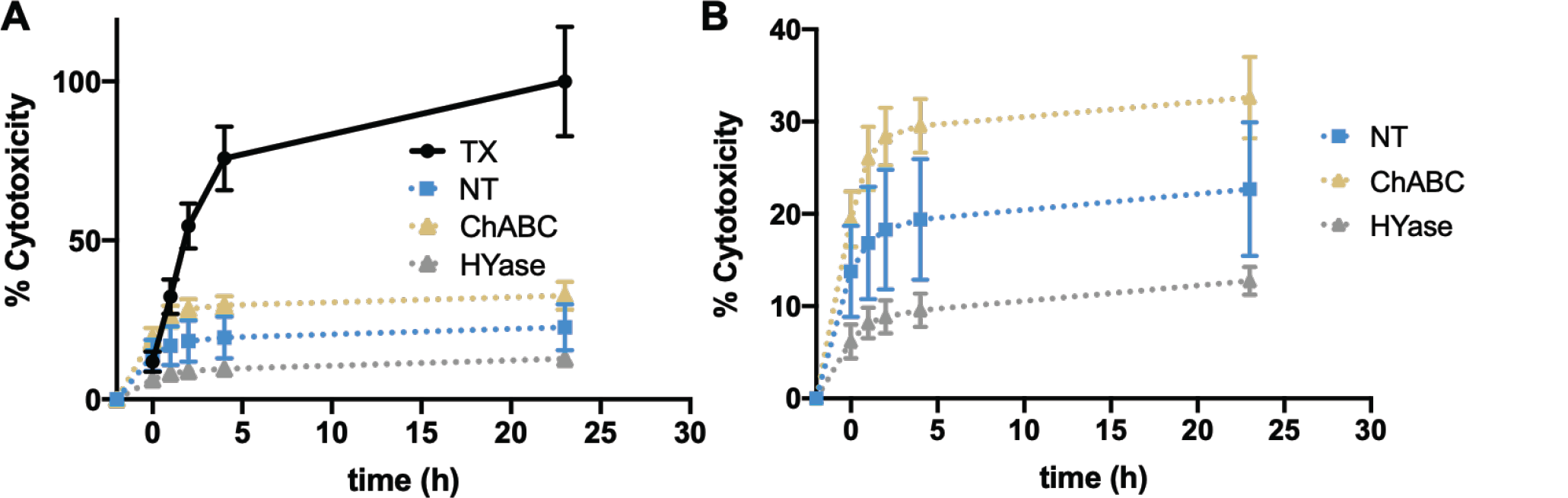
Quantifying brain slice viability following treatment with either HYase, ChABC, or SCM. (A) The LDH assay was used to quantify the % cytotoxicity versus time for brain slices treated with either Triton X-100 (black), SCM (NT, blue), Hyase (gold), or ChABC (grey). All values were normalized to the final Triton X-100 measurement. (B) Y-axis range was adjusted to better visualize the NT (blue), Hyase (gold), and ChABC (grey) groups. In all instances, scale bars represent the standard deviation of n=3 brain slices. Neither treatment (ChABC, HYase) led to a significant difference in % cytotoxicity compared to the NT control (Brown-Forsythe and Welch ANOVA test with Dunnett T3 correction).

**Table S1.**
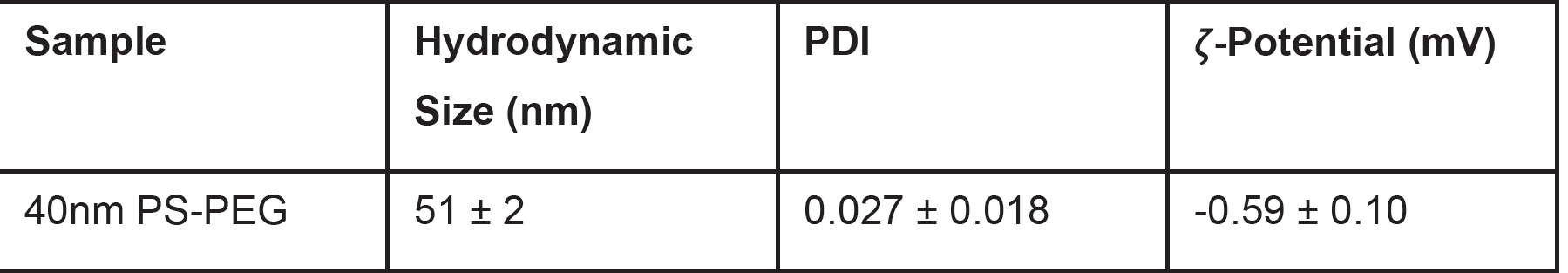
Physicochemical properties of the 40nm PS-PEG NPs. Nanoparticle hydrodynamic diameter and PDI as determined by DLS. Laser Doppler anemometry was used to determine nanoparticle ζ-potential. All experiments were performed at 25°C in 10mM NaCl, pH 7.0. Values represent the average ± standard deviation of n=3 measurements.

| Sample | Hydrodynamic Size (nm) | PDI | ζ-Potential (mV) |
| --- | --- | --- | --- |
| 40nm PS-PEG | 51 ± 2 | 0.027 ± 0.018 | -0.59 ± 0.10 |

**Table S2.**
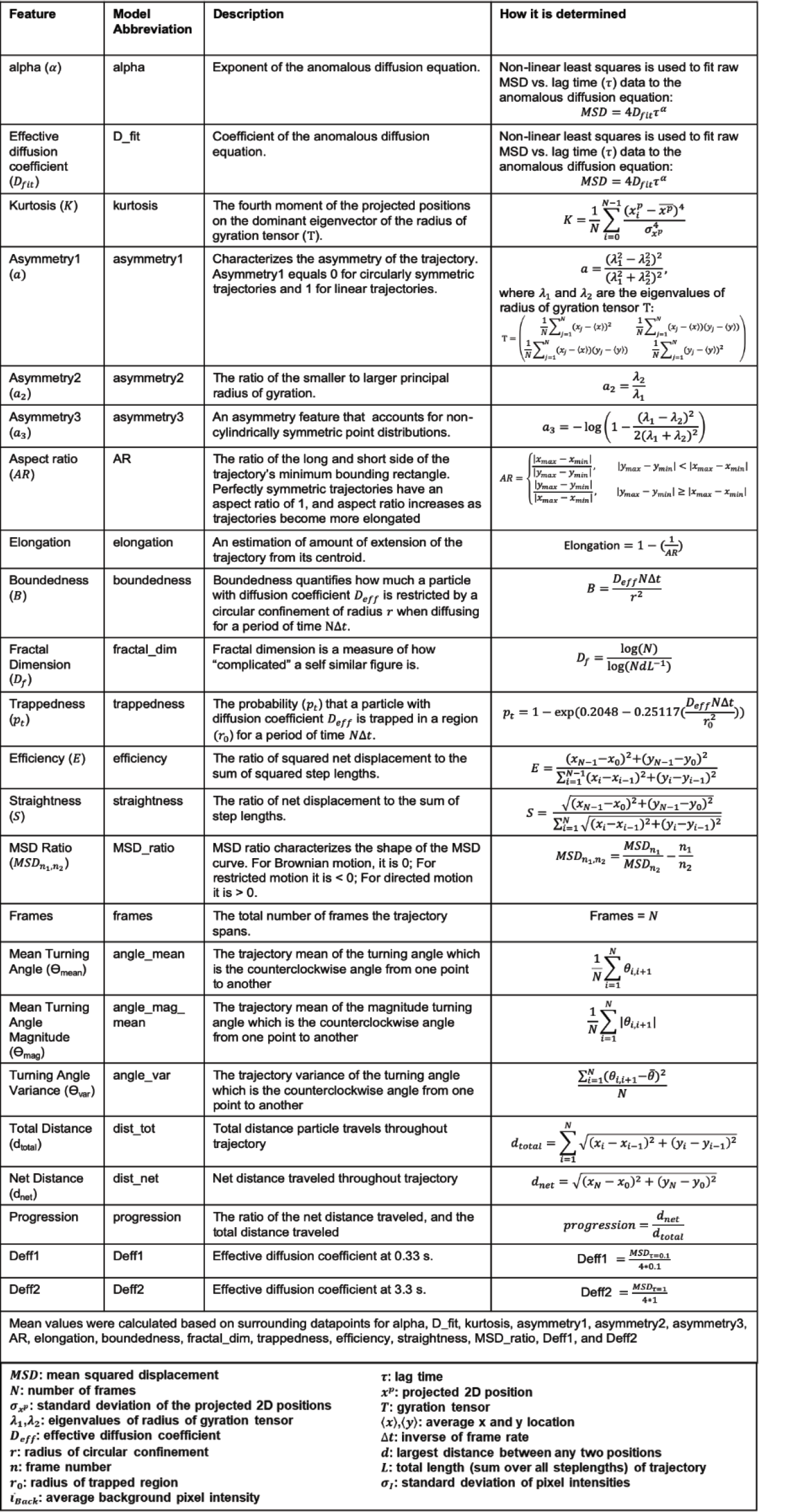
A complete list of all 39 trajectory features calculated by the diff-classifier Python package. Included for each feature is a brief description and how it is determined. Additional documentation can be found in the TraJ GitHub repository (https://github.com/thorstenwagner/TraJ.git).

**Figure S2.**
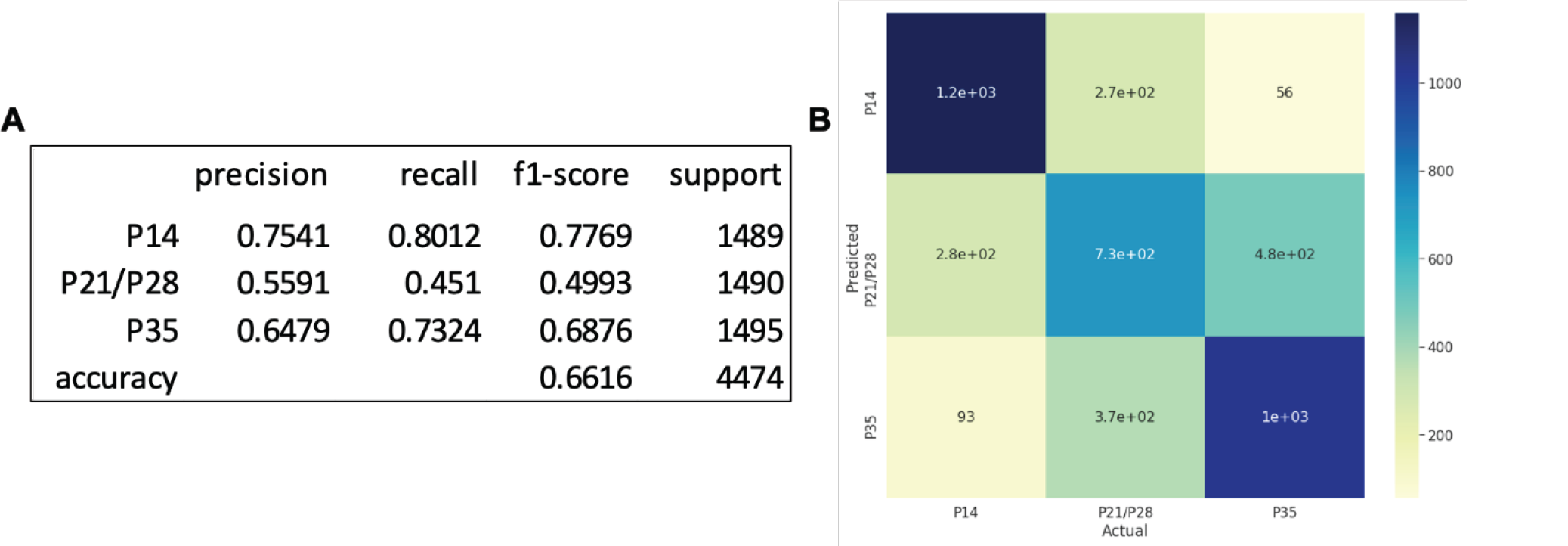
Results of XGBoost analysis for reduced resolution age-dependent data. (A) Evaluation metrics for the XGBoost classifier carried out on P14, P21/P28 combined, and P35 groups. Included are precision, recall, f1-score, and support for each age, as well as the total accuracy and support. (B) A confusion matrix displaying how predicted outcomes compare to actual classes.

**Figure S3.**
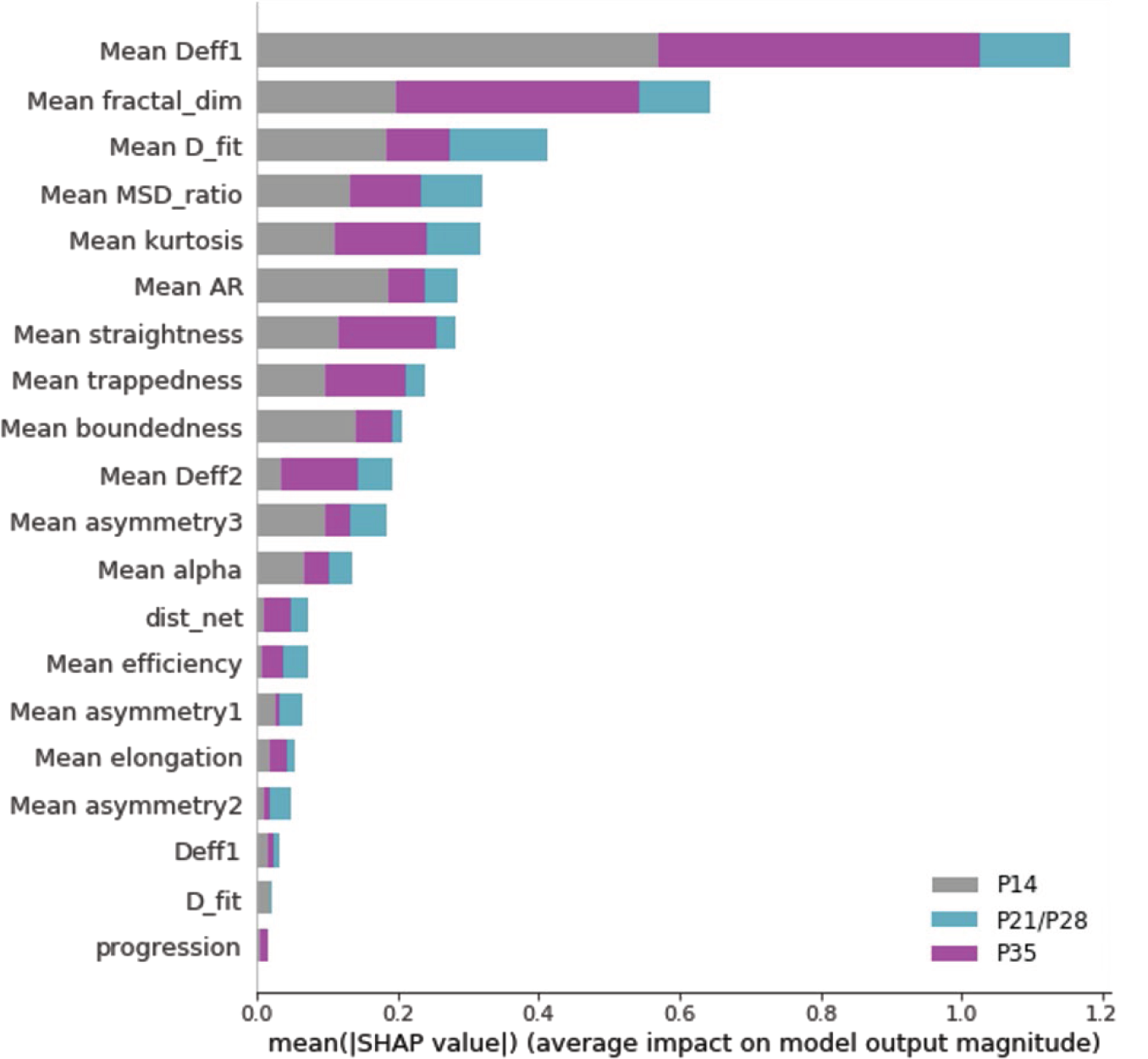
SHAP summary plot of features used in reduced resolution XGBoost classifier. Provided are the 20 most influential features. Feature importance bars are color coded to provide an indication of their importance in predicting specific age groups (grey = P14, turquoise = P21/P28 combined, purple = P35).

